# Genomic analyses identify 15 susceptibility loci and reveal *HDAC2*, *SOX2-OT*, and *IGF2BP2* in a naturally-occurring canine model of gastric cancer

**DOI:** 10.1101/2024.08.14.604426

**Authors:** Shawna R. Cook, Sanne Hugen, Jessica J. Hayward, Thomas R. Famula, Janelle M. Belanger, Elizabeth McNiel, Hille Fieten, Anita M. Oberbauer, Peter A.J. Leegwater, Elaine A. Ostrander, Paul J.J. Mandigers, Jacquelyn M. Evans

## Abstract

Gastric cancer (GC) is the fifth most common human cancer worldwide, but the genetic etiology is largely unknown. We performed a Bayesian genome-wide association study and selection analyses in a naturally-occurring canine model of GC, the Belgian Tervuren and Sheepdog breeds, to elucidate underlying genetic risk factors. We identified 15 loci with over 90% predictive accuracy for the GC phenotype. Variant filtering revealed germline putative regulatory variants for the *EPAS1* (*HIF2A*) and *PTEN* genes and a coding variant in *CD101*. Although closely related to Tervuren and Sheepdogs, Belgian Malinois rarely develop GC. Across-breed analyses uncovered protective haplotypes under selection in Malinois at *SOX2-OT* and *IGF2BP2*. Among Tervuren and Sheepdogs, *HDAC2* putative regulatory variants were present at comparatively high frequency and were associated with GC. Here, we describe a complex genetic architecture governing GC in a dog model, including genes such as *PDZRN3*, that have not been associated with human GC.

## INTRODUCTION

Gastric cancer (GC) is the fifth most common malignancy and fifth leading cause of cancer deaths in humans globally.^1^ Treatment options for GC are limited and typically include gastrectomy and chemotherapy, with a 5-year age-adjusted survival rate of 20-40%.^2–5^ Gastric adenocarcinoma comprises approximately 90% of all GCs^5^ with non-cardia involvement in >80%.^6^ The two major histological subtypes are intestinal and diffuse, based on Lauren classification.^7^ A stepwise progression from chronic inflammation through metaplasia and dysplasia characterizes the intestinal subtype.^8,9^ *Helicobacter pylori* infection is a strong environmental risk factor for intestinal GC in humans.^10^ The genetic basis of familial intestinal GC cases is poorly understood and likely polygenic.^11^ The diffuse subtype is highly infiltrative and has a worse prognosis.^12^ Diffuse GC may be caused by germline hereditary mutations in *CDH1*, encoding E-cadherin, or *CTNNA1*, which encodes part of the adherens junction protein complex.^13,14^ Approximately 10-20% of GCs show familial aggregation,^15^ but a genetic cause has only been identified for 1-3% of GCs.^16^ Human GC genome-wide association studies (GWASs) have identified common polymorphisms associated with sporadic cases of GC in some populations.^17–19^ However, the genetic risk factors underlying most cases are unknown.^15^

Gastric adenocarcinoma represents less than 1% of all neoplasias in dogs but disproportionately affects the Belgian Tervuren and Belgian Sheepdog breeds, indicating a genetic predisposition. In the Dutch Tervuren and Sheepdog populations, GC incidence is very high at 5-8% (unpublished data), and predisposition in these two breeds has been described in multiple countries spanning many years.^20–24^ GC in the Tervuren and Sheepdog is clinically and histopathologically similar to human gastric adenocarcinoma, impacting the lesser curvature of the stomach.^25,26^ Both intestinal and diffuse GC occur at approximately equal rates in Tervuren and Sheepdogs.^26^ Associations with *Helicobacter* species have not been identified in dogs.^25,27^ Consistent with the middle aged to older onset observed in humans, GC age at diagnosis is commonly 8-10 years old in these breeds.^25,28^ Humans are typically asymptomatic in early tumor stages, hindering early diagnosis and curative treatment.^29^ In dogs, initial presentation of clinical signs also occurs at advanced disease and, like humans, includes inappetence, weight loss, and vomiting.^25^ Definitive diagnosis is made by histological evaluation of endoscopic tissue biopsies.^26,30^ Surgery, with or without adjuvant chemotherapy, is the only potential curative treatment. However, even with surgery, the prognosis is very poor, with a median survival time of 178 days in dogs.^31^ Given the late stage of diagnosis and poor treatment outcomes, humane euthanasia is often elected for dogs.

The genetic structure of purebred dogs, shaped by population bottlenecks, founder events, and popular sire effects, has resulted in high within-breed homogeneity and, in some cases, enrichment for deleterious alleles.^32–34^ These factors facilitate detection of disease alleles compared to the confounding effects of heterogeneous populations in human genetic studies,^35,36^ with far fewer dogs needed than in human studies to identify high impact variants underlying complex traits.^37–39^ Causal genes and variants are often shared between dogs and humans,^40–44^ and dogs are a proven model for discovery of novel genes in human disorders.^45–47^ The Sheepdogs (also known as Groenendael) and Tervuren are members of the closely related Belgian shepherd dog group, which consists of four breeds, or varieties, based on coat type and color who share founder individuals. The most popular variety is the Belgian Malinois,^48^ which is notably not at increased risk for GC.^20–24^ The contrasting frequencies of spontaneous GC in Belgian shepherd breeds and their shared genetic background make these dogs an ideal model to elucidate the genetic basis of GC.

The aim of this study was to identify loci and candidate variants underlying GC susceptibility in Tervuren and Sheepdogs. We hypothesized that multiple loci at variable frequencies would contribute to GC risk, and that Belgian Malinois may possess a protective haplotype(s) that is at, or approaching, fixation within the breed. Applying Bayesian mixture model GWAS and cross-population number of segregating sites by length (XP-nSL) analyses to a Tervuren and Sheepdog cohort of 200 cases vs. 270 controls, we identified 15 loci that have approximately 92% predictive accuracy for GC phenotype. Across-breed selection analyses in an independent cohort of Tervuren and Sheepdogs compared to Malinois revealed additional risk and protective haplotypes.

## RESULTS

### SNP array imputation and study populations

SNP array data were imputed for 426 Belgian Tervuren, 259 Belgian Sheepdogs, and 185 Belgian Malinois, using a multi-breed reference panel of high coverage whole genome sequence (WGS) from 1,143 canines, including 20 Tervuren, 13 Sheepdogs, and 8 Malinois (Table S1). Genotype concordance was assessed by comparing imputed datasets between individuals included on multiple SNP array platforms (See Methods). Average concordance per chromosome was >85% for all allele frequency bins. The average concordance per chromosome increased to >98% for variants with INFO score ≥0.9 (n=42,223,441); these variants were used for subsequent analyses (Figure S1).

GC phenotypes were known for 200 cases (151 Tervuren, 49 Sheepdog) and 270 controls (183 Tervuren, 87 Sheepdog; Table S2). Median age at GC diagnosis or most recent health update was 9.2 and 13.2 years for cases and controls, respectively (Figure 1A). No significant association of sex with phenotype was observed (*p*=0.22). Phenotypes were unavailable for the remaining 400 samples, and this cohort was used in the selection scan analyses.

**Figure 1.**
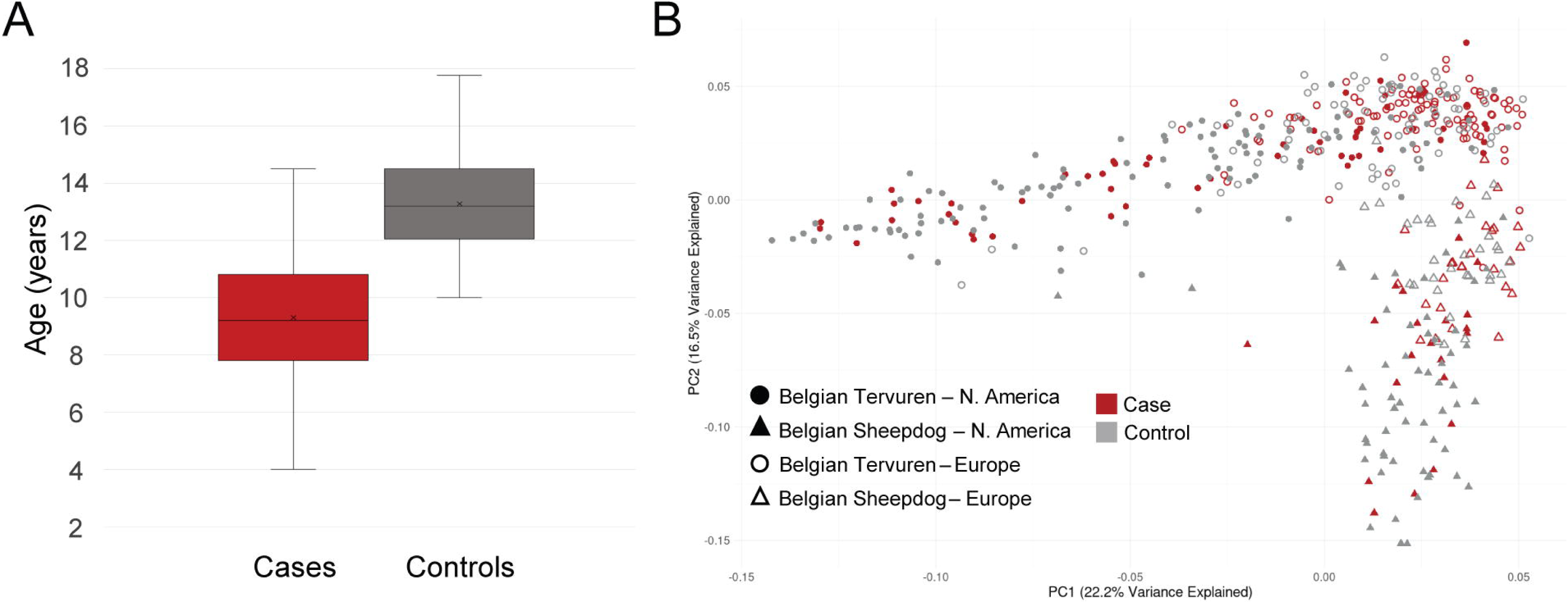
Population structure of the case-control Tervuren and Sheepdog cohort. A) Box-and-whisker plots of age at diagnosis for cases and age at collection or most recent health update for controls. B) Principal component analysis (PCA) plot for the 470 dogs included in the case vs. control analyses with each point representing an individual.

Principal component analysis (PCA) of cases and controls revealed population separation by breed/variety, as well as partial overlap between samples of European (119 cases and 105 controls) and North American (81 cases and 165 controls) origin on the first two principal components (Figure 1B). Eleven dogs reported as Tervuren or Sheepdog clustered with the opposite breed. Segregation of coat color alleles in these breeds allows fawn or mahogany Tervuren to produce solid black puppies, who may be reported as Sheepdogs, and vice versa.^49^ Breed classifications were based on PCA clusters for subsequent analyses.

### Bayesian GWAS and selection scan analysis identify multiple associated loci

Bayesian mixture model GWAS was performed for 200 cases and 270 controls, using 576,754 variants after minor allele frequency and LD pruning. Bayesian mixture modeling via BayesR^50^ simultaneously estimates the effect size of all variants rather than individual markers as in traditional GWASs, thus this approach has greater power for identifying segregating genetic risk factors for complex traits.^51–53^ Eight independent associated loci classified as having a moderate (effect size ≥0.001) effect on GC phenotype were identified across six autosomes (Figure 2A, Table 1). The locus of greatest effect size was on CFA20 and spanned only the *PDZRN3* gene. On CFA12, a 2Mb gene-dense region in high LD (r^2^>0.8) included major histocompatibility complex (MHC) class II and some class I genes. Three independent signals surpassed moderate effect on CFA26 and were proximal to the *PCDH15* and *PTEN* genes (Figure 2B). Additional signals approaching moderate effect were identified on CFA10, CFA11, and CFA24 (Table 1, Table S3).

**Figure 2.**
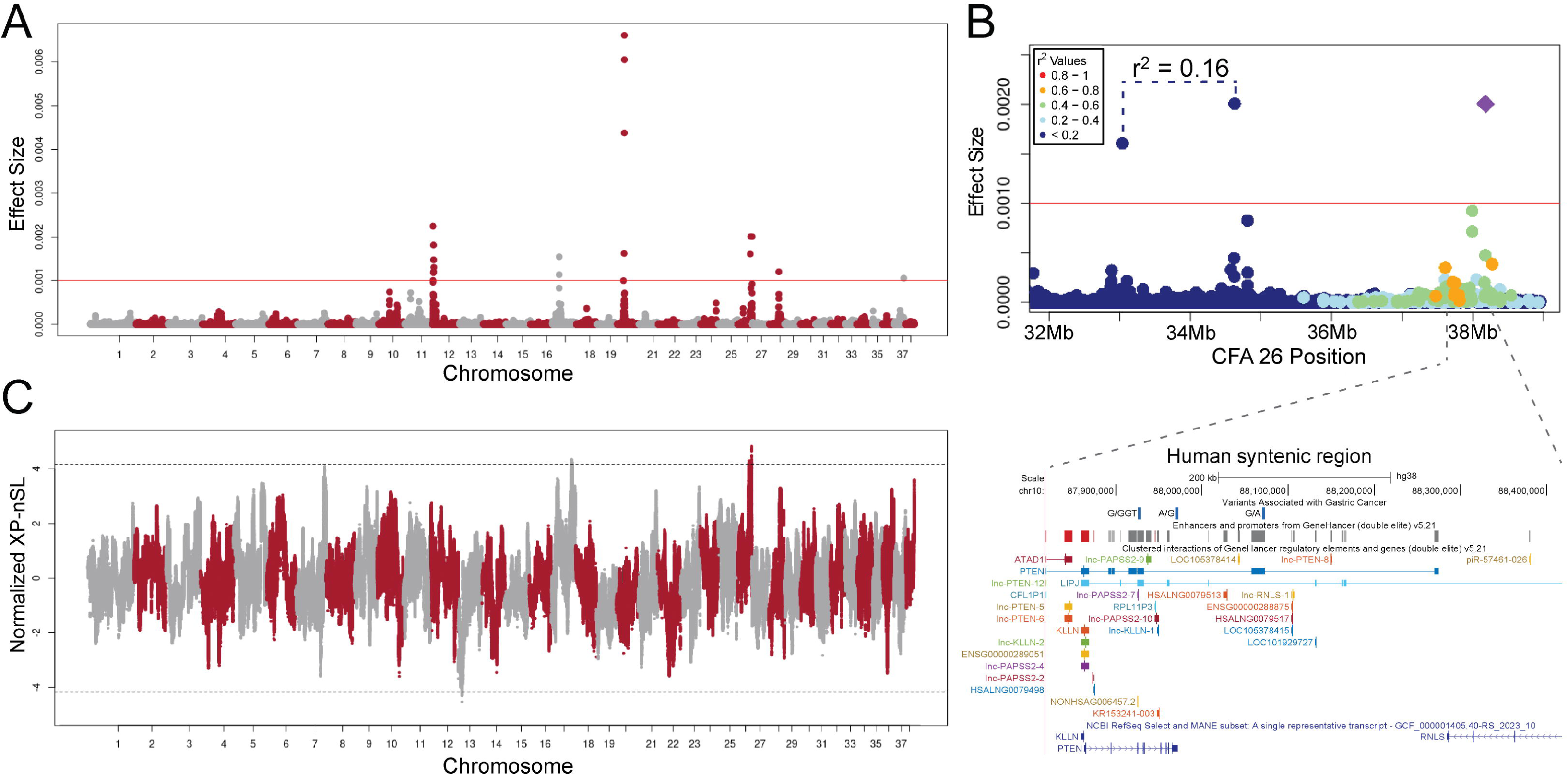
Bayesian GWAS and selection scan analysis identify multiple loci associated with gastric cancer. A) Manhattan plot showing the results of Bayesian GWAS with 200 cases vs. 270 controls. The absolute value of individual variant effect sizes (y-axis) reported by BayesR for 576,754 variants after minor allele frequency and LD pruning is plotted by position (x-axis). The red line is at the moderate effect size (0.001) threshold. B) Top: A regional Manhattan plot of CFA26 shows three independent regions surpassing moderate effect. Pairwise linkage disequilibrium (r^2^) between the top BayesR variant at CFA26:38Mb (purple diamond) and all other variants is color-coded. Bottom: UCSC image of the human (hg38) syntenic region for the CFA26:38Mb XP-nSL interval that overlaps the GWAS signal. Candidate variants meeting filtering criteria are indicated as blue vertical bars in the top track. GeneHancer double elite v5.21 enhancer (gray bars) and promoter (red bars) regulatory elements are below, with darkest colors indicating highest confidence. Clustered interactions of regulatory elements with protein coding genes and a lncRNA are shown; thick bars correspond to the regulatory element that interacts with the gene. The NCBI RefSeq Select and MANE gene track is at the bottom. C) Normalized XP-nSL values are plotted for 2,039,330 variants with dashed black lines demarcating the 99.99th percentiles. Positive values represent regions with extended haplotype homozygosity among cases, whereas negative values indicate extended haplotype homozygosity among controls.

**Table 1.** Lead variants from Bayesian mixture model GWAS and XP-nSL of 200 case and 270 control Tervuren and Sheepdogs.

| Lead Variant Position (CanFam3.1) | Critical Interval<sup>a</sup> | Analysis | BayesR Effect Size | Normalized XP-nSL | Genes within critical intervals<sup>b</sup> |
| --- | --- | --- | --- | --- | --- |
| CFA10:28,537,583 | CFA10:28,152,671-28,821,673 | GWAS | 0.000741 | - | *RBFOX2, APOL5, APOL6, MB, RASD2, MCM5, HMOX1, TOM1* |
| CFA10:48,471,044 | CFA10:48,141,927-48,512,574 | GWAS | 0.000442 | - | *PRKCE* |
| CFA11:15,714,902 | CFA11:15,377,666-15,728,753 | GWAS | 0.000723 | - | *GRAMD2B* |
| CFA11:38,106,700 | CFA11:38,104,905-38,206,955 | GWAS | 0.000521 | - | *ADAMTSL1* |
| CFA12:1,359,325 | CFA12:1,359,325-3,451,568 | GWAS | 0.00225 | - | 115 genes and non-coding transcripts |
| CFA13:9,923,659 | CFA13:9,831,529-10,417,265 | XP-nSL | - | -4.53 | *NUDCD1, ENY2, PKHD1L1, EBAG9, SYBU, SYBU-AS1* |
| CFA17:18,030,682 | CFA17:17,946,446-18,030,682 | GWAS | 0.00154 | - | *KLHL29* |
| CFA17:54,006,088 | CFA17:53,974,863-54,445,506 | XP-nSL | - | 4.35 | *CD58, IGSF3, CD2, PTGFRN, CD101, CD101-AS1* |
| CFA17:55,324,675 | CFA17:55,013,848-55,324,755 | XP-nSL | - | 4.25 | *GDAP2, WDR3, SPAG17* |
| CFA20:18,681,361 | CFA20:18,646,458-18,718,242 | GWAS | 0.00661 | - | *PDZRN3* |
| CFA24:41,167,902 | CFA24:41,030,826-41,191,757 | GWAS | 0.000484 | - | - |
| CFA26:31,661,685 | CFA26:31,661,685-32,050,496 | XP-nSL | - | 4.30 | - |
| CFA26:33,035,867 | CFA26:32,881,257-33,106,991 | GWAS | 0.00161 | - | *PCDH15* |
| CFA26:34,625,869 | CFA26:34,612,341-34,848,102 | GWAS | 0.00201 | - | - |
| CFA26:38,012,570 | CFA26:37,794,344-38,275,926 | XP-nSL | - | 4.83 | *ATAD1, CFL1P1, KLLN, PTEN, RNLS* |
| CFA26:38,180,502 | CFA26:38,180,502 | GWAS | 0.00200 | - | *RNLS* |
| CFA28:24,778,275 | CFA28:24,295,247-24,847,272 | GWAS | 0.00120 | - | *HABP2, NRAP, CASP7, PLEKHS1, DCLRE1A, NHLRC2* |
| CFA37:26,075,098 | CFA37:26,075,098 | GWAS | 0.00105 | - | *OBSL1* |

<sup>a</sup>GWAS critical intervals were defined by variants in high LD with the lead SNV ( $r^2 \geq 0.8$ ). XP-nSL critical intervals represent regions in the top 0.01%.
<sup>b</sup>All genes within the critical interval, in CanFam3.1 or CanFam4, are reported with the exception of CFA12 because it contains 115 genes.

XP-nSL analysis^54^ was used to compare cases and controls to identify regions of increased homozygosity and haplotype length in either group (Figure 2C, Table 1). The strongest signal replicated the CFA26:38Mb association from the GWAS at *PTEN*. An additional selection signal was identified proximal to the CFA26:32Mb GWAS signal. These regions demonstrated extended haplotype homozygosity among cases and are associated with GC risk (Table 1, Table S4). Regions within the 99.99th percentile (Table S4) also included loci on CFA13:9.9Mb, CFA17:54Mb, and CFA17:55.3Mb, demonstrating increased homozygosity among controls (CFA13) or cases (CFA17; Table 1). Gene ontology of the CFA17 regions indicated enrichment for immunoglobulin genes (FDR *p*=0.024), including *IGSF3*, *CD2*, *CD58*, and *CD101*.

### Two loci are associated with intestinal gastric cancer subtype

For a subset of the cases, the diffuse (n=25) vs. intestinal (n=26) histopathological tumor subtype was known.^26^ Fisher’s exact tests were performed at the lead variant/haplotype for each of the aforementioned loci to detect significant associations with tumor subtype. The CFA17:54Mb (Fisher’s 2-tailed *p*=0.016, OR=3.61, 95% CI=1.28-10.18) and CFA28:24.8Mb (Fisher’s 2-tailed *p*=0.028, OR=2.62, 95% CI=1.17-5.86) loci were each associated with intestinal tumor subtype (Table S5). Significant associations with the diffuse subtype were not detected.

### Fifteen loci predict gastric cancer phenotype with greater than 90% discriminatory accuracy

Bayesian GWAS and XP-nSL case-control analyses identified 15 independent loci (See Methods). At these loci, the total number of risk alleles/haplotypes (2n=30) observed among cases averaged 20.3 compared to 15.9 among controls (two-tailed *t*-test *p*=1.31x10^-61^). Malinois (n=149), who are at low risk for GC, possessed an average of 15.4 risk alleles, clustering with the controls (Figure 3A). In the imputation reference panel, representing 227 other breeds, the average number of risk alleles per individual (n=1,102) was 13.6, which is within the range observed in controls (Table S6). Age at GC diagnosis was negatively correlated with total number of risk alleles (*p*=0.00184), *i.e.*, dogs possessing high numbers of risk alleles were diagnosed with GC at younger ages (Figure S2).

**Figure 3.**
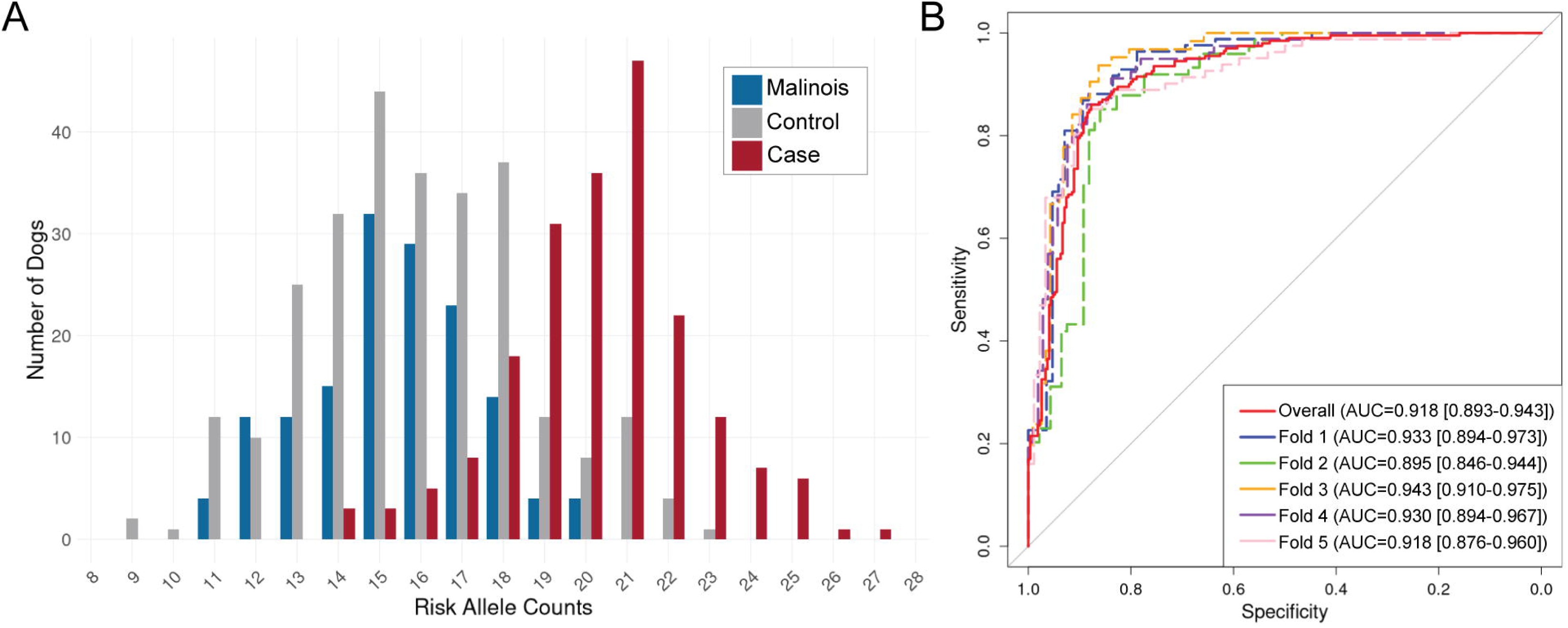
Fifteen loci predict gastric cancer phenotype with greater than 90% discriminatory accuracy. A) Bar graph of the distribution of risk alleles among cases (n=200, red), controls (n=270, gray), and Malinois (n=149, blue). Cases have an average of 20.3 total risk alleles while controls have 15.9 (*p*=1.31x10^-61^), and Malinois cluster with controls. B) Receiver Operating Characteristics (ROC) curves for the 5-fold cross-validated 15-locus additive model of GC risk in Tervuren and Sheepdog. Area Under the Curve (AUC) is reported for each fold, with the 95% confidence interval in brackets.

To identify the best model for predicting GC phenotype, additive models including top GWAS variants and three-SNP haplotypes of case vs. control XP-nSL loci were compared using ANOVA Chi-squared tests (Table S7). Significant evidence of epistatic interactions between the 15 loci was not observed, thus additive models weighting the effects of each locus were applied, rather than interactive models. A 13-locus model including the eight loci of moderate effect and five loci approaching moderate effect threshold from the GWAS was a significant improvement over the eight moderate loci alone (*p*=4.32x10^-11^). Adding two independent loci identified through XP-nSL significantly improved GC phenotype prediction (*p*=0.025), thus including the top 15 loci from the genomic analyses produces the simplest and most accurate predictive model. Five-fold cross-validation of this 15-locus additive model within the GWAS case-control cohort yielded an ROC-AUC of 0.918 (95% CI=0.893-0.943), indicating that this model predicts GC phenotype with greater than 90% discriminatory accuracy (Figure 3B). CFA12 and CFA20 genotypes were the most influential predictors of phenotype in the model (Figure S3).

### Belgian shepherd across-breed comparisons identify GC risk and protective loci under selection

We hypothesized that some loci under selection in Malinois may confer protection from developing GC, based on the breed’s low risk despite their close genetic relationship to the high-risk Tervuren and Sheepdogs.^23,24,26,55^ Further, some loci at high frequency in both Tervuren and Sheepdogs may contribute to their GC predisposition. An independent population of dogs (non-GWAS cohort) was used to identify regions at, or approaching, fixation in the breeds. After relationship pruning, 69 Tervuren, 82 Sheepdogs, and 149 Malinois were included in XP-nSL (Figure 4A, Table S8), Fst (Figure S4, Table S9), and runs of homozygosity (ROH; Figure S5, Table S10) analyses.

**Figure 4.**
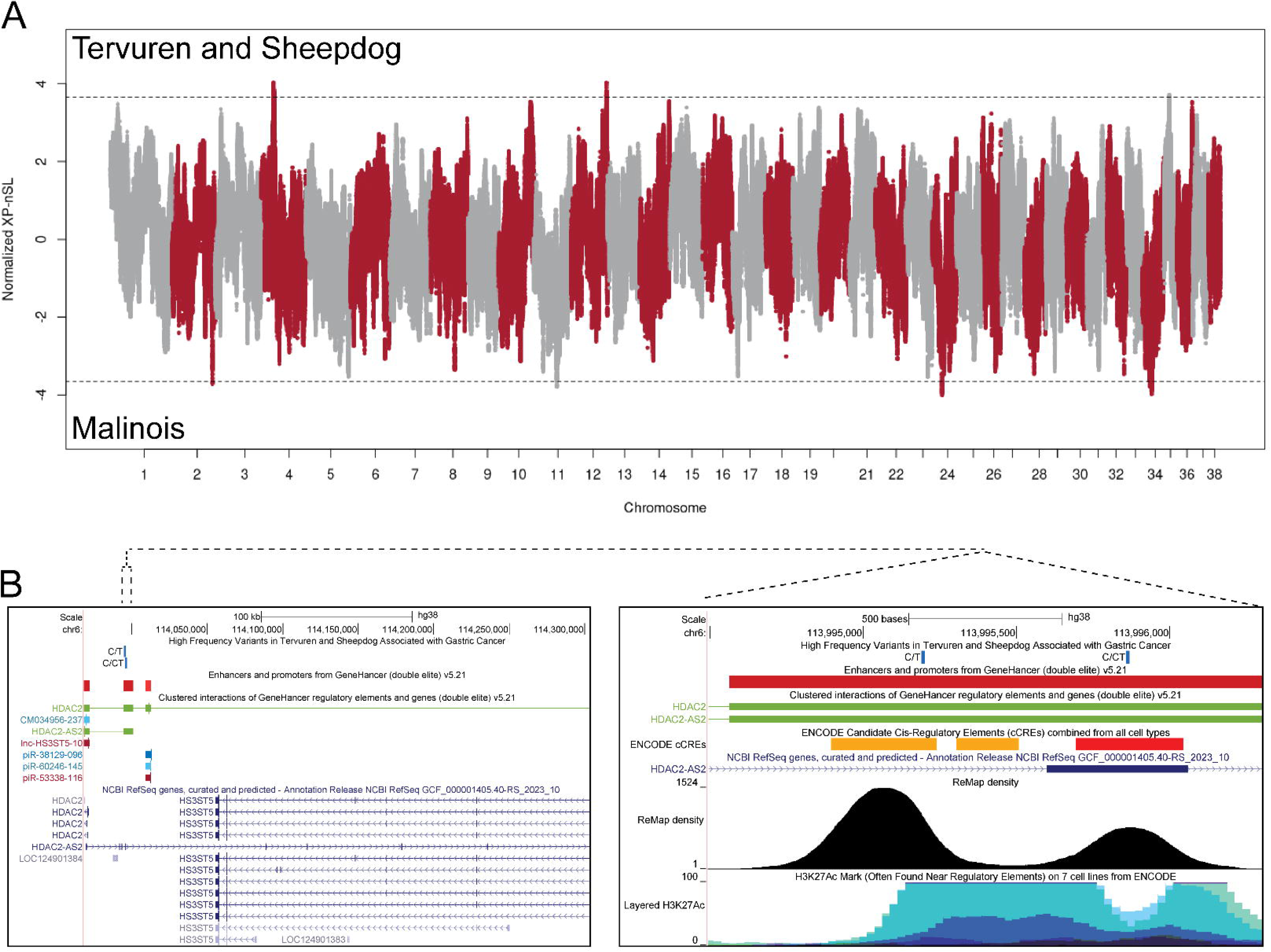
Belgian shepherd across-breed comparisons identify GC risk and protective loci under selection. A) Normalized XP-nSL values are plotted for 2,039,330 variants with dashed black lines demarcating the 99.99th percentiles. Positive values indicate regions under selection in an independent Tervuren (n=69) and Sheepdog (n=82) cohort with unknown GC phenotypes, whereas negative values represent regions under selection in Malinois (n=149). B) Left: UCSC image of the human (hg38) syntenic CFA12 region under selection in Tervuren and Sheepdogs. Two variants significantly associated with GC are denoted, as well as all GeneHancer double elite regulatory elements. Right: zooming in to the human syntenic location of the two identified variants shows overlap with a GeneHancer double elite element and ENCODE candidate cis-regulatory elements. The second variant falls within an untranslated region of *HDAC2-AS2*. Below, the ReMap ChIP-seq density track displays evidence of transcriptional regulators, and the ENCODE H3K27ac track shows enrichment of the H3K27ac histone modification, indicating areas of active enhancers.

The strongest evidence for shared selection among the Tervuren and Sheepdog was at CFA4:24.2-24.4Mb (haplotype frequency = 95.4%), with the locus observed in all three selection analyses (Table 2). The gene-rich region includes genes that are dysregulated in human GC tumors, such as *PLAU*, *FUT11*, and *VCL*. Strong selection among Tervuren and Sheepdogs was also identified by XP-nSL and Fst on CFA12:70Mb, including *HS3ST5* and the upstream region and first exon of *HDAC2* (Table 2). These regions show significant enrichment for genes involved in the biosynthesis of heparan sulfate and heparin (FDR *p*=0.023) based on *NDST2* on CFA4 and *HS3ST5* on CFA12.

**Table 2.**
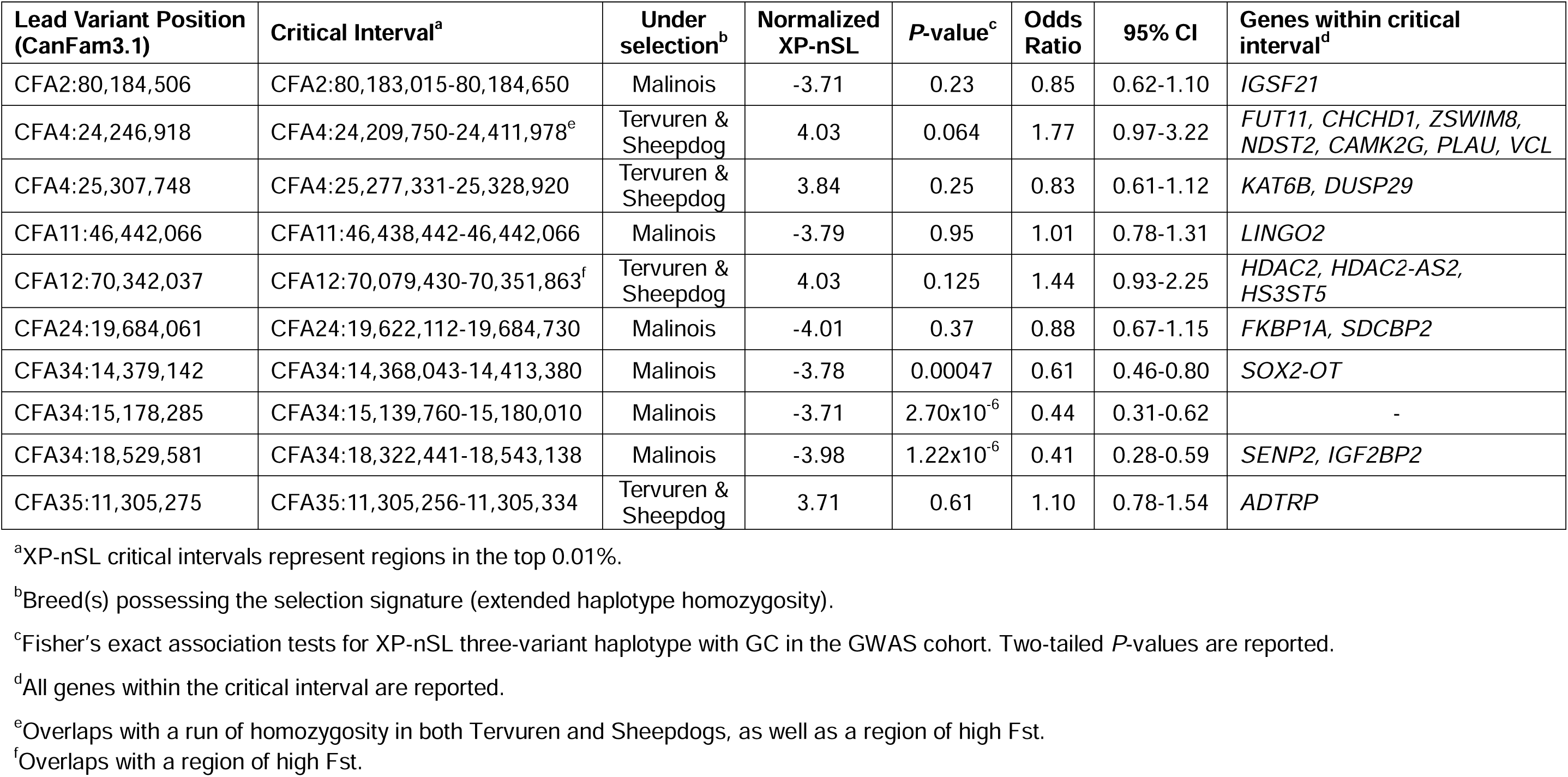
Regions under selection in Tervuren and Sheepdogs or Malinois identified through XP-nSL.

Six loci under strong selection in the Malinois were identified through XP-nSL (Figure 4A). A haplotype including *SOX2-OT* (CFA34:14.3-14.4Mb) was present among 100% of Malinois, with 91.3% haplotype frequency (Table 2). The Malinois haplotype was observed at significantly higher frequency among the GWAS control cohort (46% in controls vs. 34% in cases, Fisher’s 2-tailed *p*=0.00047, OR=0.61, 95% CI=0.47-0.80), suggesting a protective effect. A second locus under selection in Malinois at CFA34:18.3-18.5Mb, harboring *IGF2BP2*, is also overrepresented among controls (24.1% haplotype frequency in controls vs. 11.5% in cases, Fisher’s 2-tailed *p*=1.22x10^-6^, OR=0.41, 95% CI=0.28-0.59). The remaining Malinois XP-nSL haplotypes were not significantly associated with GC phenotype in the GWAS cohort. Adding the CFA4, 12, and 34 loci individually did not significantly improve upon the 15-locus GC prediction model (Table S7). The model including the CFA4 locus was slightly preferred by leave one out cross validation information criterion (looic) but was not significantly better than the base model.

### Rare variants confer risk or protection for gastric cancer

To identify candidate variants driving GC susceptibility in the Tervuren and Sheepdog, critical intervals defined through the above genomic analyses were interrogated in high-coverage WGS data for coding or regulatory variants unique to these breeds or rarely observed in other breeds (See Methods). Variants present on a risk or protective haplotype were pruned according to their effect on protein or PhyloP conservation scores and their presence in human (hg38) syntenic regulatory elements.

Two coding and nine non-coding variants were associated with GC in the case-control cohort, having *P*-values less than or equal to those of the lead variants from GWAS or selection scans (Table 3). Within the GWAS identified regions, coding variants predicted to impact protein function were present in *CD101* (XP_533019.3:p.Leu934Pro) and *DCLRE1A* (XP_535018.3:p.Leu326Arg). The former is on the risk haplotype and replicates the lead variant association with intestinal tumor subtype. The *DCLRE1A* variant was identified on a control haplotype and did not replicate the intestinal subtype association of the CFA28 lead variant. The CFA28 locus may harbor both risk and protective haplotypes relevant to GC. Variants within GWAS identified regions were present in multiple syntenic human enhancer elements for *EPAS1* (*HIF2A*) and were associated with GC (Table 3). Putative regulatory variants of *PTEN* were also identified, including within an enhancer demonstrating *HDAC2* binding,^56^ and within 3’UTR of human *PTEN* transcripts (Figure 2B, Table 3). The only lead GWAS variant to meet the allele frequency filtering criterion was on CFA12 (alternate allele AF=8.3%); however, it was filtered out at the conservation score step (PhyloP = -0.658). The variant does lie within a human syntenic enhancer (GH06J031896).

**Table 3.** Coding and putative regulatory candidate variants for gastric cancer susceptibility.

| Candidate Variant Position (CanFam3.1) | REF/ ALT Allele<sup>a</sup> | ALT AF Tervuren & Sheepdog (2n=66)<sup>b</sup> | ALT AF other canines (2n=2220) | GeneHancer Double Elite Element ID<sup>c</sup> | Element Type | Gene/Gene Target | PhyloP<sup>d</sup> | P-value Lead Variant<sup>e</sup> | P-value of candidate variant<sup>f</sup> | OR | 95% CI |
| --- | --- | --- | --- | --- | --- | --- | --- | --- | --- | --- | --- |
| CFA4:24,304,634 | G/A | 0.88 | 0.034 | GH10J073878 | Enhancer | *PLAU, FUT11* | 4.84 | 0.064 | 0.064 | 1.82 | 0.998-3.31 |
| CFA10:48,313,427 | A/G | 0.29 | 0.032 | GH02J046009 | Enhancer | *EPAS1/HIF2A* | 1.86 | 0.00015 | 2.92x10<sup>-6</sup> | 1.91 | 1.46-2.50 |
| CFA10:48,330,465 | T/C | 0.29 | 0.069 | GH02J046029 | Enhancer | *EPAS1/HIF2A* | 3.12 | 0.00015 | 2.92x10<sup>-6</sup> | 1.91 | 1.46-2.50 |
| CFA10:48,378,432 | G/T | 0.26 | 0.011 | GH02J046085 | Enhancer | *EPAS1/HIF2A* | 2.87 | 0.00015 | 2.92x10<sup>-6</sup> | 1.91 | 1.46-2.50 |
| CFA10:48,399,845 | G/T | 0.29 | 0.015 | GH02J046116 | Enhancer | *EPAS1/HIF2A* | 1.05 | 0.00015 | 2.92x10<sup>-6</sup> | 1.91 | 1.46-2.50 |
| CFA12:70,106,510 | C/T | 0.83 | 0.073 | GH06J113994 | Enhancer, Promoter | *HDAC2, HDAC2-AS2* | 1.29 | 0.125 | 0.0020 | 1.88 | 1.26-2.82 |
| CFA12:70,107,159 | C/CT | 0.79 | 0.0059 | GH06J113994 | Enhancer, Promoter | *HDAC2, HDAC2-AS2* | 1.13 | 0.125 | 0.013 | 1.60 | 1.11-2.29 |
| CFA17:54,417,724 | T/C | 0.29 | 0.012 | - | - | *CD101* | 2.85 | 7.05x10<sup>-10</sup> | 8.81x10<sup>-12</sup> | 2.51 | 1.92-3.28 |
| CFA26:37,880,356 | G/ GGT | 0.53 | 0.066 | GH10J087915 | Enhancer | *PTEN, LIPJ, Inc-PAPSS2-7* | 1.36 | 9.08x10<sup>-9</sup> | 2.08x10<sup>-10</sup> | 2.38 | 1.82-3.13 |
| CFA26:37,917,914 | A/G | 0.55 | 0.072 | - | 3'UTR<sup>g</sup> | *PTEN* | 1.58 | 9.08x10<sup>-9</sup> | 2.08x10<sup>-10</sup> | 2.38 | 1.82-3.13 |
| CFA26:37,984,898 | G/A | 0.53 | 0.086 | GH10J088058 | Enhancer | *PTEN* | 6.90 | 9.08x10<sup>-9</sup> | 5.29x10<sup>-13</sup> | 2.69 | 2.05-3.54 |
| CFA28:24,720,256 | A/C | 0.44 | 0.038 | - | - | *DCLRE1A* | 2.89 | 0.00074 | 0.00027 | 0.60 | 0.46-0.79 |

<sup>a</sup>Reference allele (REF) is based on WGS reference panel, and alternate allele (ALT) is alternate to the reference.
<sup>b</sup>Allele frequencies (AF) of the ALT allele are calculated using the WGS reference panel (n=1143).
<sup>c</sup>GeneHancer double elite v5.20 IDs for regulatory elements encompassing the human (hg38) syntenic position of the dog candidate variant.
<sup>d</sup>PhyloP scores from 241 vertebrates are based on the human (hg38) variant positions.
<sup>e</sup>Fisher's exact two-tailed *P*-value of BayesR lead variant or XP-nSL three-variant haplotype were calculated comparing GWAS cases (2n=400) to controls (2n=540).
<sup>f</sup>Fisher's exact two-tailed *P*-value of the imputed candidate variant genotypes for GWAS cases (2n=400) vs. controls (2n=540).
<sup>g</sup>This variant is syntenic to the 3'UTR of human *PTEN* transcripts per NCBI RefSeq (NM\_001304718.2, NM\_000314.8, NM\_001304717.5)

Within the CFA4 region of strong selection in Tervuren and Sheepdogs, one non-coding variant approached significantly increased frequency among GC cases (*p*=0.064). This variant is syntenic to a human super enhancer (SEdb2.0 ID: SE_02_152500349) expressed in stomach tissue and interacting with *PLAU* and *FUT11*. The CFA12:70Mb Tervuren and Sheepdog haplotype harbors two putative regulatory variants of *HDAC2* that were significantly associated with GC in the GWAS cohort (*p*=0.0020-0.013). These variants lie within a syntenic human super enhancer (SEdb2.0 ID: SE_02_019600073) and promoter element expressed in the stomach. The CFA12:70,107,159 variant lifts over to an untranslated region of human antisense-RNA (*HDAC2-AS2*) regulating *HDAC2* and *HS3ST5* (Figure 4B). Incorporating the candidate variants individually into the 15-locus model did not significantly improve upon utilizing the lead variants (Table S11).

## DISCUSSION

GC is a leading cause of cancer-related deaths in humans worldwide with poor treatment options and prognosis. The genetic etiology is largely unknown. In dogs, Belgian Tervuren and Belgian Sheepdog breeds are at a uniquely high risk of developing GC, while their close breed relative, the Belgian Malinois, is at low risk. These breeds represent a naturally occurring model to elucidate the genetic basis of GC. Here, we have conducted the first GC GWAS in dogs, uncovering 15 loci that predict the phenotype with 92% discriminatory accuracy. These include the tumor suppressor gene *PTEN*, as well as genes that have not yet been associated with human GC, such as *PDZRN3* and *KLHL29*. Leveraging the contrasting GC incidences between these breeds, we performed across-breed selection scans and uncovered an additional risk locus at *HDAC2* approaching fixation in Tervuren and Sheepdogs and two protective loci at *SOX2-OT* and *IGF2BP2* at high frequency in Malinois. These genes have critical roles in human GC progression and represent potential therapeutic targets.^57–60^

Bayesian GWAS for GC in Tervuren and Sheepdogs revealed 13 loci associated with GC susceptibility. The top variant is intronic to *PDZRN3* (*LNX3*), an E3 ubiquitin ligase with roles in angiogenesis and cell migration through non-canonical Wnt signaling pathways.^61,62^ In humans, PDZRN3 ubiquitinates DVL3,^61^ which is overexpressed in diffuse GC,^63^ and PDZRN3 is reported to regulate connexin 43 (Cx43) protein stability in gap junctions.^64^ Connexins are linked to multiple cancers,^65,66^ and Cx43 is dysregulated in GC tumors.^67–69^ PDZRN3 is a promising therapeutic target in colorectal cancer.^70^ A second locus of moderate effect harbors *KLHL29*, a member of the Kelch-like (KLHL) protein family that bind to specific substrate protein targets for ubiquitination by the cullin3 E3 ubiquitin ligase complex. KLHLs have been implicated in human gastrointestinal tumors,^71^ including *KLHL5*, *KLHL9*, and *KLHL19*/*KEAP1* in GC.^72–74^ A target of KLHL29, DDX3, regulates E-cadherin,^75–78^ which is often inactivated in diffuse GC,^79^ and germline variants in *CDH1*, which encodes E-cadherin, are associated with hereditary diffuse GC in humans.^13,14^

Three independent GWAS signals were detected on CFA26, including a locus at *PTEN*, which was also identified in the case-control XP-nSL analysis. *PTEN* is a well-known tumor suppressor gene whose loss has been identified in many tumors, including human GC.^80,81^ Decreased expression of *PTEN* has also been reported in the gastric mucosa of first-degree relatives to GC patients.^82^ We identified variants syntenic to two human enhancers that interact with *PTEN*, one of which is known to be expressed in the stomach, and a 3’UTR variant. Additional putative regulatory variants were identified within the CFA10 GWAS critical interval syntenic to multiple human enhancers of *EPAS1*, also known as *HIF2A*. *EPAS1* expression is induced under hypoxia, and it is upregulated in many malignancies, including GC where it contributes to metastasis.^83–90^

The CFA12 locus has a strong effect on GC phenotype prediction. This gene-dense region harbors major histocompatibility complex class II, as well as some of the class I genes. Alleles of MHC class II HLA genes have been associated with risk or protection from human GC,^91,92^ as well as autoimmune gastritis,^93,94^ which can progress to GC.^95^ The relationship between gastritis and GC in dogs is unclear.^25^ Further work is necessary to identify the specific variants at this locus contributing to GC predisposition in Tervuren and Sheepdogs. Additional associations with immune-related genes were identified through our case vs. control selection analyses. A CFA13 region of extended homozygosity among controls contains *EBAG9*, also known as tumor associated antigen *RCAS1*, which is involved in tumor immune evasion and demonstrates aberrant localized expression in human GC tumors.^96–99^ There may be a recessive protective variant in this region or a dominant risk factor present among cases, which were more heterozygous in this region. Enrichment of immunoglobulin genes was observed within an extended case haplotype on CFA17:54Mb. Variant filtering revealed a rare protein coding variant within *CD101* that disrupts a conserved amino acid, is enriched among cases, and is associated with intestinal tumor subtype in our cohort. *CD101* has roles in inhibiting T cell proliferation, promoting development of regulatory T cells, and controlling inflammation, and germline *CD101* coding variants have been shown to have a proinflammatory effect in humans.^100–102^

Within dog breeds, many loci are under strong artificial selection for breed-defining physical and behavioral traits, such as body size, coat color and type, intelligence, and trainability.^103^ As a result, deleterious alleles can increase in frequency through hitchhiking or pleiotropic effects.^43,104,105^ We performed across-breed analyses to identify regions under divergent selection in the high-risk Tervuren and Sheepdog compared to the low-risk Malinois. The close genetic relatedness of these breeds^55^ minimized the number of loci resulting from this approach, highlighting a subset of genomic regions contributing to GC susceptibility. In Malinois, a haplotype including the lncRNA *SOX2-OT* is under strong selection and is associated with protection from GC. A SNP intronic to *SOX2-OT* was associated with decreased risk of developing GC in humans.^106^ *SOX2-OT* dysregulation occurs in various malignancies,^107^ including GC where it is overexpressed, leading to epithelial-mesenchymal transition.^57^ A key role of *SOX2-OT* is modulating expression of the *SOX2* gene.^108^ *SOX2* possesses antitumor effects in GC and demonstrates positive regulation of *PTEN*.^109,110^ A gradual reduction in *SOX2* expression has been observed during GC tumorigenesis.^109^ In addition to critical roles in the stomach,^111,112^ *SOX2-OT* and *SOX2* are important for neural differentiation,^108,113,114^ and this locus may be under selection in Malinois for cognitive traits.

A second protective locus was identified in Malinois at *IGF2BP2*. In addition to roles in metabolism and type 2 diabetes, *IGF2BP2* contributes to cancer proliferation and cancer stem cell maintenance.^115^ Overexpression of *IGF2BP2* promotes GC progression and epithelial to mesenchymal transition in human GC tumors and cell lines,^58,116,117^ and SNPs have been associated with chemotherapeutic response in GC.^118^ IGF2BP2 is a recognition binding protein for N6-methyladenosine (m^6^A) RNA methylation, involved in mRNA stability, and its interaction with multiple targets promotes human GC tumor progression and represents a potential therapeutic target.^58,60,119,120^ The use of small molecule inhibitors of IGF2BP2 in other cancers have shown promising anti-cancer effects.^121,122^

Among Tervuren and Sheepdogs, a shared haplotype on CFA4 is under strong selection and harbors variants within human syntenic enhancer elements of *PLAU* and *FUT11* that are skewed toward GC cases (*p*=0.064). *PLAU* is a driver of tumor progression in multiple cancers and is overexpressed in GC.^123,124^ *FUT11* is a fucosyltransferase that is overexpressed in GC and a predictor of poor prognosis.^125,126^ Gene amplification and missense mutations of *FUT11* are commonly observed in GC tumors.^126^ This region also contains *NDST2*, which plays a role in modification of heparan sulfate chains via deacetylation and sulfation of glucosamine.^127^

A second shared haplotype under strong selection in Tervuren and Sheepdogs is on CFA12 and includes *HS3ST5* and the upstream region and first exon of *HDAC2*. Like *NDST2*, *HS3ST5* is in the heparan sulfate metabolism pathway. In GC, heparan sulfate proteoglycan levels decrease with tumor progression,^128^ and dysregulation of other heparan sulfate modifying genes is associated with GC progression and prognosis.^129^ However, the specific role of *HS3ST5* in GC is unclear.^130^ *HDAC2* is a class I histone deacetylase that has roles in transcriptional regulation, cell proliferation, and tumorigenesis.^59^ We identified putative regulatory variants, syntenic to a human super enhancer, that are associated with GC and are rarely found outside of the Tervuren and Sheepdog breeds. *HDAC2* is upregulated in human GC tumors, and knockdown reduces cell proliferation and restores *CDKN2A* expression in GC cell lines.^131–135^ HDAC2 interacts with CDH1 (E-cadherin), complete loss of which causes diffuse GC.^136^ HDAC inhibitors have been shown to have anti-proliferative effects in GC cell lines and are a potential therapy for GC.^135^

Multiple genes identified herein have reported gene/protein interactions (Figure S6), including downregulation of *PTEN* by *HDAC2* and formation of HDAC2-SOX2 protein complexes in cancer stemness.^59^ While dysregulation of many of these genes has been described in human GC, this work supports the need for functional investigation of their interactions in GC tumors, which may facilitate development of targeted therapies.

### Limitations of the study

The five-fold cross validation approach applied herein indicates 92% predictive accuracy of the model; however, additional samples will be necessary to assess model accuracy in an independent validation cohort to ensure the model is not overfitting. This study does not account for environmental risk factors for GC. Population substructure was present based on geographic origin and breed; however, we applied a relationship matrix in the GWAS to correct for this. Statistical evidence of genotype interactions was not observed. This is likely due to limitations in sample size to robustly represent all possible combinations of genotypes across the 15 loci. Because GC is rare across dog breeds, we applied a variant filtering approach that identified candidate variants at low frequency outside of Tervuren and Sheepdogs. However, there may be common polymorphisms at one or more risk loci that rarely exist in combination in other breeds. Further studies will be necessary to assess relevant common polymorphisms. Next steps also include functional validation of putative regulatory variants identified herein. While this work highlights shared genetic risk factors for GC in two high-risk breeds, future studies should include definitive Lauren tumor subtypes to determine whether additional loci distinguish predisposition to intestinal vs. diffuse GC.

### Conclusions

This is the first study in dogs to identify genetic variants associated with GC, revealing 15 loci predictive of GC phenotype. The shared genetic background and contrasting incidences of GC among high-risk Tervuren and Sheepdogs compared to low-risk Malinois enabled identification of additional genetic risk or protective factors at high frequency in either population. This work establishes a genetic risk model for these breeds that will enable selective breeding to reduce disease frequency, as well as routine monitoring for high-risk individuals. Genes identified herein overlap with those dysregulated in human GC tumors and include previously unassociated genes, such as *PDZRN3* and *KLHL29*. Here we demonstrate that GC in Tervuren and Sheepdogs reflects the genetic complexity observed in humans, in addition to clinical heterogeneity, and thus has tremendous translational potential to impact human health.

## MATERIALS & METHODS

### Samples

Whole blood or buccal samples were obtained from purebred Belgian Shepherd breeds with informed owner consent in accordance with regulations of each institution’s animal care and use committee. DNA was isolated according to the Puregene DNA Isolation protocol (Gentra) or standard phenol-chloroform extraction. Cases were diagnosed via clinical signs and ultrasound, histopathology, and/or cytology. Approximately 84% of cases had diagnoses confirmed by histopathology. For inclusion as controls, dogs were at least 10 years of age at sample collection or last health update with no owner-reported history of GC or clinical signs. A total of 224 samples were collected from Europe (119 cases and 105 controls) and 246 samples from North America (81 cases and 165 controls). Additional samples and/or SNP array genotype data were collected irrespective of health status and age at collection for breed-based population comparisons (Belgian Tervuren, n=95; Belgian Sheepdog, n=120; Belgian Malinois, n=185).

### Whole genome sequence reference panel

Whole genome sequence data were generated for 20 Belgian Tervuren (9 cases, 8 controls, and 3 of unknown phenotype) and 13 Belgian Sheepdogs (7 cases, 4 controls, and 2 of unknown phenotype), ranging from 9.6 to 48.2X coverage, with a median coverage of 16X. Raw data were aligned to CanFam3.1, and these genomes were included in a multibreed joint-called reference panel comprising 1,143 canines representing 230 breeds, including 8 Belgian Malinois and 1 Belgian Laekenois (Table S1). Alignment to CanFam3.1 and jointcalling were performed as described by Plassais et al. (2019).^137^

Phasing was performed for each chromosome, using SHAPEIT5^138^ phase_common command in 30Mb chunks with 10Mb overlapping, followed by ligation with SHAPEIT5 ligate command. The output was converted to VCF and indexed using BCFtools^139^ prior to input into imp5Converter to create an IMPUTE5^140^ file for use in the imputation pipeline.

### Pre-imputation SNP array concordance and quality control

SNP array genotypes were generated on either the Illumina CanineHD BeadChip array (173k markers), Embark’s custom high-density Illumina 220k SNP arrays,^141^ or Affymetrix Axiom Canine HD Array (670k markers; Table S2). Each array dataset (target panel) was imputed independently prior to merging all three datasets. First, the conform-gt tool was used to unify allele strand between the target and reference panels and to remove variants that were only present in the target panel.^142^ Fifteen dogs were genotyped on both the 173k and 220k arrays, 23 dogs on the 173k and 670k arrays, and 7 dogs on the 220k and 670k arrays. Three of these dogs were genotyped on all three arrays. To identify discordant SNPs between duplicate dogs on the arrays, we generated transposed (--export A-transpose command using PLINK v2.00a3.7LM (24 Oct 2022)) files for each chromosome of each array dataset, where the genotypes were coded 0, 1, 2 for homozygous alternate, heterozygous, and homozygous reference, respectively.^143^ The transposed files were used as input in the python script “imputation_accuracy.py,” which compares all sites in common across every dataset type for all duplicate dogs. The script removes all sites with allele mismatches. The output is a matrix of values calculated by subtracting the genotype codes (0,1,2) in one dataset from the other. A value of 0 for a given site means that the genotypes for the two dogs in the two datasets are the same. A site that was discordant in any of the comparisons was removed from all three genotype datasets. These matrix outputs were used in the python script “analysisOfTSV.py” to calculate concordance across sites in common for each chromosome per dog, and then concordance across dogs in common per site. A site that was discordant in any of the genotype comparisons in at least one of the duplicate dogs was removed from all genotype datasets, resulting in 5,300 markers removed from 173k, 5,518 markers removed from 220k, and 5,602 markers removed from the 670k.

### SNP array imputation

Genotype datasets were filtered in PLINK 1.9^143^ to remove SNPs with MAF <0.01 and/or call rates <90%, and then split into chromosomes. PLINK files were then converted into VCF files in VcfCooker v1.1.1 (https://genome.sph.umich.edu/wiki/VcfCooker) then indexed using BCFtools index. Phasing was performed using SHAPEIT5 v5.1.0 phase_common command incorporating available parent-offspring information (four parent-offspring pairs and one trio for 173k; one parent and five offspring for 220k) and genetic map files for the 173k and 220k datasets. Default SHAPEIT5 options were used with the exception of pbwt-depth, which was set to eight, to improve accuracy while not compromising speed. For the imputation step, imp5Chunker, part of the IMPUTE5 package, was first used to split the chromosomes into ∼5-10Mb chunks with ∼500kb buffer region, which were used to impute across using the reference panel (--h option in IMPUTE5 v1.1.5). The imputed chunks were ligated back together using BCFtools concat. Finally, PLINK2 was used to generate pgen format PLINK files.

### Imputation concordance

To calculate imputation concordance between dogs included in multiple datasets, we used the same scripts as above. The imputed pgen output files were converted into .traw files and fed into “imputation_accuracy.py” and the output matrix files were fed into “analysisOfTSV.py.” Concordance values for all variants shared across the imputed output files were calculated in non-reference allele frequency bins of 0.05 increments. After imputation and merging, variants with INFO score ≥ 0.9 (Table S12), variants missing in at least one array dataset (n=17,222,525) were removed using PLINK1.9,^144^ resulting in 42,223,441 remaining variants.

### Genome-wide association analyses

Pruning for minor allele frequency <0.05 in the GC GWAS cohort yielded 5,998,047 variants. For PCA, variants were pruned for linkage disequilibrium (r^2^>0.4) using the --indep-pairwise option with 50 kb windows in PLINK1.9, resulting in 130,872 variants. PCA was conducted using eigensoft’s smartpca.^145^ For GWAS, a genomic relatedness matrix was created in GEMMA^146^ using the LD-pruned dataset of 130,872 variants. The eigenvectors were subsequently extracted and used as covariates for GWAS analyses to correct for population substructure. BayesR^50^ was first used to determine effect sizes of the 5,998,047 variants, using 300,000 iterations, 100,000 burn-in, permutations, and GEMMA eigenvector covariates. The overall estimated effect each SNV has on the phenotype is updated iteratively. In each iteration, BayesR estimates the probability of an individual variant belonging to one of four normal distributions: zero, low, moderate, and high effect, each explaining 0%, 0.01%, 0.1%, and 1% of the genetic variance, respectively. Critical intervals were defined by the most centromeric and telomeric variants in pairwise LD (r^2^ ≥ 0.8) with the lead variant in each independent GWAS peak. To obtain an accurate estimation of effect sizes at each locus, which are diluted by variants in strong LD,^50,147^ LD-based clumping in PLINK1.9^144^ was used to filter variants with r^2^ ≥ 0.9 relative to the lead variant within 50 kb windows, while maintaining the lead variant of greatest effect size. In total, 576,754 variants were retained, and BayesR analysis was repeated using the aforementioned parameters.

### Genotype models

For regions identified in GWAS and XP-nSL that overlapped or were in high LD, a single representative variant or haplotype was used for subsequent analyses, specifically the CFA17:54,006,088 three-SNP haplotype, CFA26:33,035,867 variant, and CFA26:38,180,502 variant (Table 1). Preference was given to GWAS lead variants. The case predominant allele or haplotype was assigned as “risk,” and the total number of risk alleles/haplotypes present in each individual in the case-control population was determined. A two-tailed *t*-test was used to calculate the difference between the risk allele load of cases and controls. The total risk allele/haplotype count in the Belgian Malinois (n=149) was calculated using the same methodology. Lastly, the total risk allele count was calculated across canines in the imputation reference panel, excluding Tervuren, Sheepdogs, and Malinois (n=1102).

Pairwise interactions between BayesR and XP-nSL lead variants were analyzed using the “--epistasis” command in PLINK1.9,^144^ and multiple testing correction was applied to determine significance. Generalized linear models were created in R^148^ to determine the model that best explains GC risk in the GWAS cohort. These models were compared via ANOVA chi-square testing. Additionally, models were generated using a Bayesian generalized linear model via the stan_glm command in the rstanarm package^149^ and evaluated using leave-one-out cross-validation information criterion (LOOIC) performed using 25,000 iterations, 5,000 burn-in, and a thinning parameter of 25.

K-fold cross-validation was used to evaluate the additive model which incorporated the 15 loci identified from the GWAS and case vs. control XP-nSL analyses. The GWAS cohort was randomly partitioned into five subsets. An iterative approach was applied wherein four subsets were used to train the model and the remaining subset was used to test the model. The receiver operating characteristic curve and area under the curve (ROC-AUC) were calculated using the pROC package in R for each of the five iterations and the average. Variable importance of each locus was calculated using the caret package in R to determine the loci having the greatest influence within the model.^150^

### Selection scans

#### Cross-population number of segregating sites by length (XP-nSL)

Two populations of Belgian Shepherd dogs, the GWAS cohort (n=470) and an independent cohort with unknown GC phenotypes (n=401), were examined for evidence of selective sweeps that may contribute to overall GC risk. For the independent cohort with unknown GC phenotypes, dogs were split by breed (n=185 Malinois, 120 Sheepdogs, and 95 Tervuren), and a minor allele frequency of 0.1 was applied prior to genetic relatedness pruning with GCTA grm-cutoff^151^ of 0.3, resulting in 149 Malinois, 82 Sheepdogs, and 69 Tervuren for the across-breed analysis.

The initial genotyping dataset comprising 42,223,441 variants was LD-pruned using a pairwise r^2^ of 0.99 and a sliding window of 50 kb, with 2,039,330 variants remaining for analysis. SelScan default settings were used to perform XP-nSL followed by “norm”^54,152^ to identify regions under selection in cases or controls in the GWAS cohort, as well as to compare Tervuren and Sheepdogs to the Malinois in the cohort of unknown GC phenotypes. Variants within the top 99.99% threshold were further examined and used to identify critical intervals. Haplotype frequencies were calculated using the lead variant and two flanking variants reported by XP-nSL. Fisher’s exact association tests were performed in the case-control GWAS cohort comparing the frequency of the predominant haplotype in the population under selection to the frequency of all other haplotypes.

#### Runs of Homozygosity & Fst

Runs of homozygosity (ROH) were assessed separately in the Tervuren, Sheepdogs, and Malinois in the cohort of dogs with unknown GC phenotype. The LD-pruned dataset of 2,039,330 variants was evaluated using the sliding window method in detectRuns.^153^ The sliding window used 40 variants, of which one was allowed to be heterozygous, and a minimum ROH length was 500kb. As the data were imputed and sufficiently dense, minimum density and maximum gap size parameters were set to be ignored. The default window threshold value of 0.05 was used. Regions of interest were defined per breed as the genomic areas that were most commonly shared within that breed, specifically, those in the 99.9th percentile.

Fst was calculated in PLINK1.9^144^ using the same population of dogs, comparing the Tervuren and Sheepdogs to the Malinois. For this analysis, the initial dataset of 42,223,441 variants was used and monomorphic variants were discarded (n=7,836,277 variants remaining). The top 0.01% of Fst values were extracted and variants within 1Mb were collapsed into one region.

### Variant filtration

All variants present in the WGS reference dataset of 1,143 canids (n=33 Tervuren and Sheepdog) within the critical intervals identified through GWAS were extracted. Additionally, all variants within the XP-nSL critical intervals +/-100kb buffer on either side were extracted. Variants were filtered per individual for genotype quality score ≥20 via BCFtools. SNPs and INDELs with alternate allele frequency (AF) ≥ 0.25 in Tervuren and Sheepdogs and alternate AF ≤ 0.1 in all other canids were retained. To allow for the reference allele being a candidate GC variant, filtering was also performed to retain variants with alternate AF ≤0.75 in Tervuren and Sheepdogs and alternate AF ≥0.9 in other canids. Variant effect predictor (VEP) was used to annotate variants.^154^ PhyloP scores from 241 vertebrates^155^ were extracted, and variants with scores >1 were retained. Putative regulatory variants were identified from syntenic human regulatory elements using the hg38 GeneHancer v5.20 track and evidence of super enhancers was examined in SEdb 2.0.^156^ SNPs and INDELs meeting the above criteria were manually examined in relation to all imputed variants within the critical region to determine if the identified variants segregated with a risk or non-risk haplotype as identified in the GWAS cohort. Fisher’s exact two-tailed *p*-values were calculated for candidate variants in the imputed GWAS dataset (2n=940) to determine if allele frequencies were significantly different between cases and controls. Allele frequencies (Table 3) are reported for the entire reference population (2n=2286).

### Enrichment analysis

All gene set enrichment analyses were conducted in STRING version 12.0.^157^ Regions of interest were converted to their equivalent human positions using UCSC’s LiftOver^158^ and input into STRING using *Homo sapiens* as the organism.

## DATA AVAILABILITY

The python scripts “imputation_accuracy.py” and “analysisOfTSV.py” used in the concordance part of the imputation pipeline are publicly available on github at: https://github.com/JessicaHayward/imputation_accuracy_GC. SNP array and imputed data will be available at datadryad.org upon publication. SRA accession numbers for high coverage WGS are available in Table S1.

## ACKNOWLEDGEMENTS

We thank the many owners, breeders, and veterinarians who contributed samples and clinical data for this study. We also thank Embark Veterinary, Inc. for providing anonymized array data. This material is based upon work supported by the Cornell Richard P. Riney Canine Health Center Research Grants Program, a grant made available to the College of Veterinary Medicine, Cornell University. This work was supported by grants from the American Kennel Club Canine Health Foundation (AMO), the Intramural Program of the National Human Genome Research Institute at the National Institutes of Health (EAO), The Hartwell Foundation (SRC), Albert C. Bostwick Foundation, Belgian Sheepdog Club of America, American Belgian Tervuren Club, Nederlandse Vereniging van Belgische Herdershonden, Belgische Herder Club Nederland, Deutscher Klub für Belgische Schäferhunde, the Tjech KCHBO, and the Nederlands Kankerfonds voor Dieren.

## AUTHOR CONTRIBUTIONS

Writing-original draft -JME, SRC, SH, JJH

Writing-review & editing - JME, SRC, EAO, AMO, JMB, JJH, HF, SH, PJJM, PAJL

Investigation & Formal analysis - SRC, JME, SH, JJH, TRF

Data curation - JJH, SRC, SH, JMB, JME Validation - SRC, TRF, JME

Visualization - SRC, JJH, JME

Resources - EAO, AMO, JMB, EM, HF, PAJL, PJJM, JME

Conceptualization - JME, SRC, EAO, AMO, PAJL, PJJM

Funding acquisition - JME, EAO, PJJM, SRC, AMO

## DECLARATION OF INTERESTS

The authors declare no competing interests.

## SUPPLEMENTARY FIGURES & TABLES

Table S1. WGS imputation reference panel SRA numbers.

Table S2. Sample information for all imputed Belgian Tervuren, Sheepdog, and Malinois data.

Table S3. BayesR GWAS results for all variants with effect size ≥ 0.0001.

Table S4. XP-nSL 99.99th percentile results of case vs. control selection scan.

Table S5. Fisher’s exact tests for association of each case-control lead variant or haplotype with tumor subtype.

Table S6. Total number of risk alleles across the 15 loci included in GC risk model for each individual in the imputation reference panel.

Table S7. Results of model comparisons using ANOVA Chi-squared tests and leave one out cross validation information criterion (LOOIC).

Table S8. XP-nSL 99.99th percentile results of Belgian Tervuren and Sheepdog vs. Belgian Malinois selection scan.

Table S9. Top 99.99th percentile Fst values calculated in PLINK1.9 with imputed data for Belgian Tervuren and Belgian Sheepdogs vs. Belgian Malinois.

Table S10. Regions in the 99.9th percentile of per breed runs of homozygosity for Belgian Tervuren, Belgian Sheepdog, and Belgian Malinois breeds.

Table S11. Results of model comparisons using identified variants using ANOVA Chi-squared tests.

Table S12. Number of variants in each SNP array dataset after imputation, INFO score pruning, and merging.

**Figure S1. Imputation concordance between dogs genotyped on multiple arrays.** A) Average concordance per chromosome of all imputed variants prior to INFO score filtering. B) Average concordance after removing variants with INFO score <0.9. Each datapoint is the average genotype concordance per alternate allele frequency bin per chromosome. Concordance for dogs genotyped on the 173k and 220k arrays (n=15) is shown in red, 173k and 670k (n=23) in green, and 220k and 670k (n=7) in blue.

**Figure S2. Linear model demonstrating the relationship between number of risk alleles and age at gastric cancer diagnosis.** The total number of risk alleles, among cases, in the 15-locus model (2n=30) is plotted on the x-axis, with age at diagnosis in years on the y-axis.

**Figure S3. Importance of number of risk alleles at each locus in the 15-locus predictive model.** The importance score (x-axis) ranks the relative contribution of one (1) or two (2) risk alleles at each locus (y-axis) to the model’s predictive accuracy, with the highest score indicating the greatest effect within the model. The importance score is based on the absolute value of the t-statistic of each variable when applied to the model and was calculated using the varImp function in the caret R package. Importance is set to zero for zero copies of the risk allele.

**Figure S4. Across-breed fixation index (Fst) values.** A) Fst values of 7,836,277 non-monomorphic variants comparing the Belgian Tervuren (n=69) and Belgian Sheepdogs (n=82) of unknown phenotype to Belgian Malinois (n=149) as reported by PLINK1.9. The dashed black line represents the 99.99% threshold. B) Distribution of Fst values; approximately half of all values lie between 0 and 0.05.

**Figure S5. Incidence plots of runs of homozygosity (ROH) by breed.** ROH were assessed in the Tervuren (n=69), Sheepdog (n=82), and Malinois (n=149) for 2,039,330 variants, with a minimum 500kb interval. The proportion of times each SNV is represented in an ROH within the population is shown. The dashed line represents the 99.9th percentile of runs calculated per breed.

**Figure S6. STRING protein network interactions.** Proteins identified through GWAS and XP-nSL genomic analyses with reported interactions were included, as well as CDH1, whose expression is often lost in human diffuse GC. Protein interactions as defined by default STRING options are represented by gray lines. Blue shapes indicate downregulation and red shapes indicate upregulation of the protein or gene in human GC, while gray represents unknown. The figure was generated with Cytoscape.

## Notes

### Competing Interest Statement

The authors have declared no competing interest.

