## Supplemental Figures for "Genomic analyses identify 15 susceptibility loci and reveal *HDAC2*, *SOX2-OT*, and *IGF2BP2* in a naturally-occurring canine model of gastric cancer"

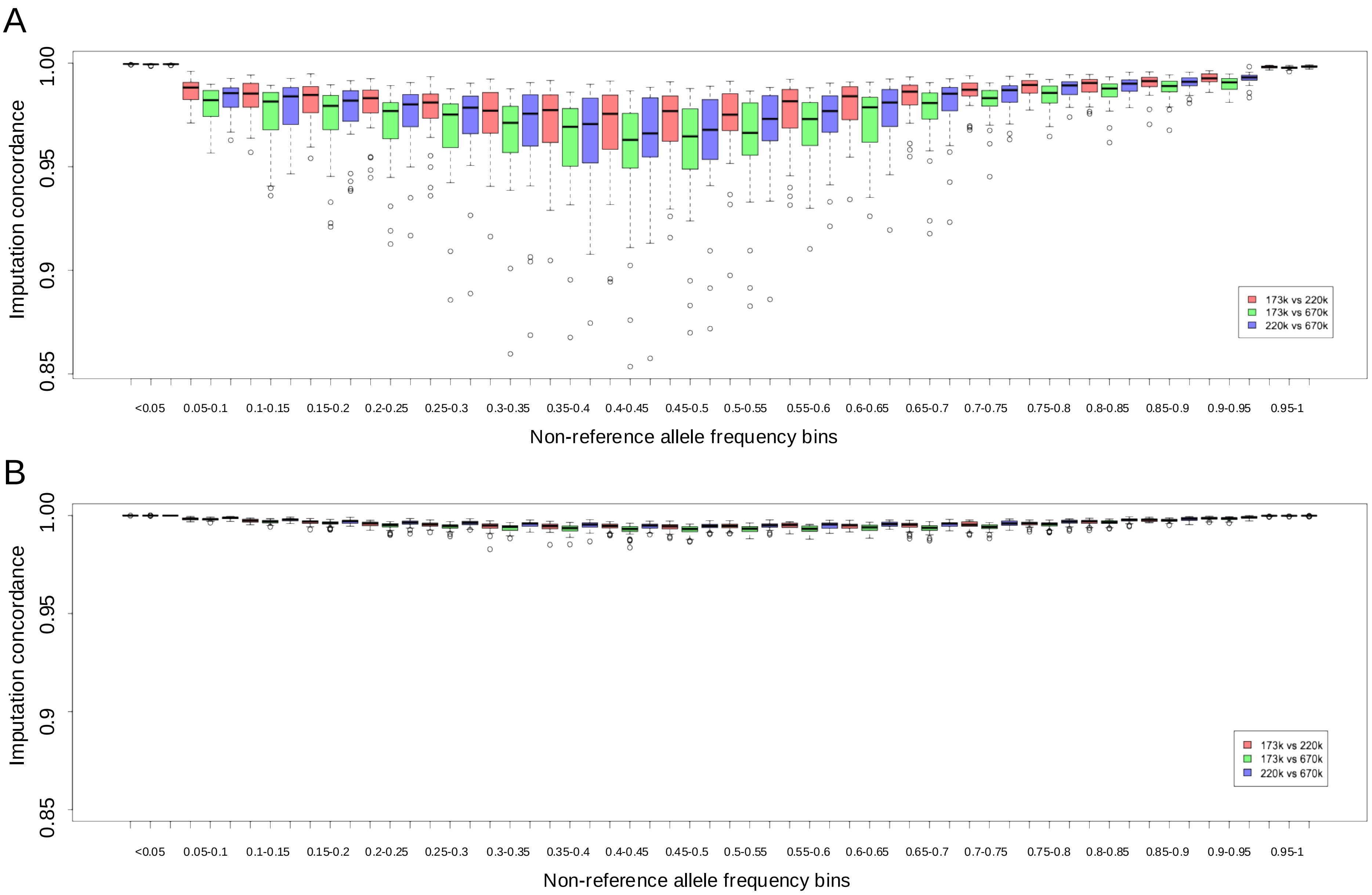


Figure S1. Imputation concordance between dogs genotyped on multiple arrays. A) Average concordance per chromosome of all imputed variants prior to INFO score filtering. B) Average concordance after removing variants with INFO score <0.9. Each datapoint is the average genotype concordance per alternate allele frequency bin per chromosome. Concordance for dogs genotyped on the 173k and 220k arrays (n=15) is shown in red, 173k and 670k (n=23) in green, and 220k and 670k (n=7) in blue.

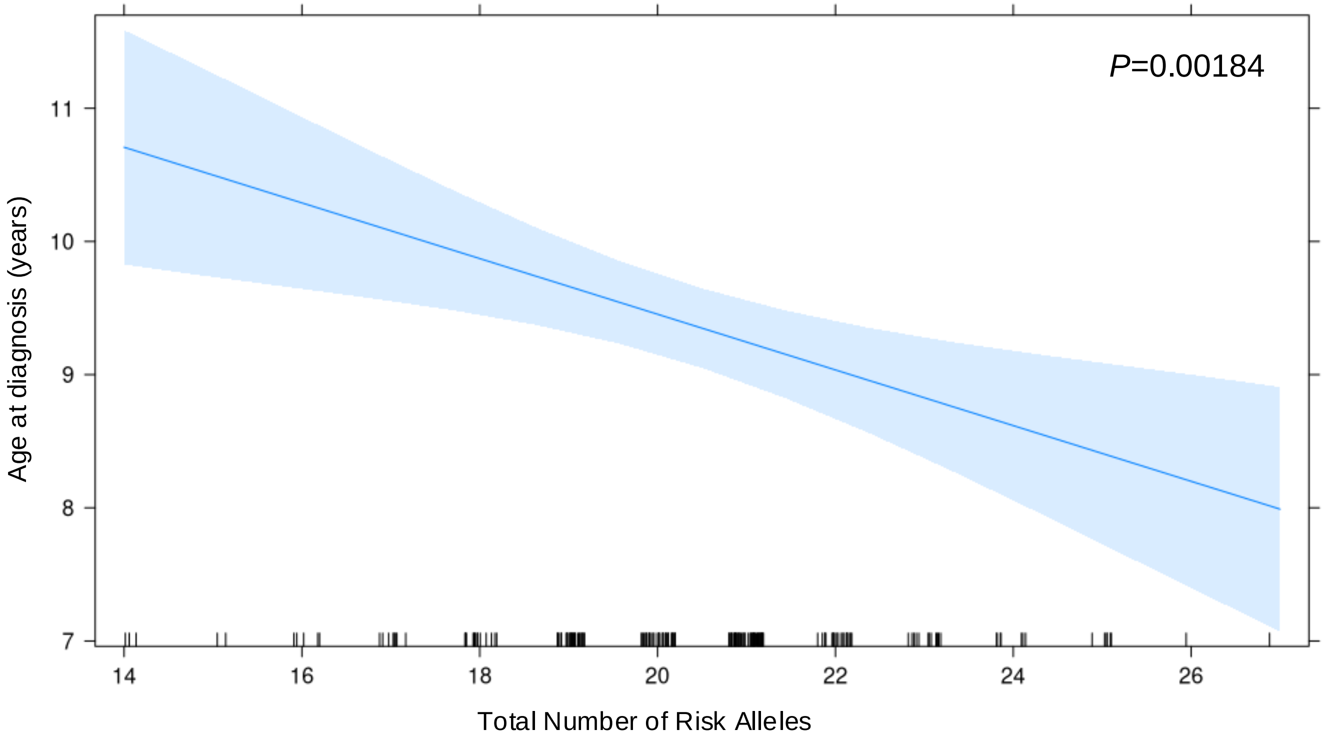


Figure S2. Linear model demonstrating the relationship between number of risk alleles and age at gastric cancer diagnosis. The total number of risk alleles, among cases, in the 15-locus model (2n=30) is plotted on the x-axis, with age at diagnosis in years on the y-axis.

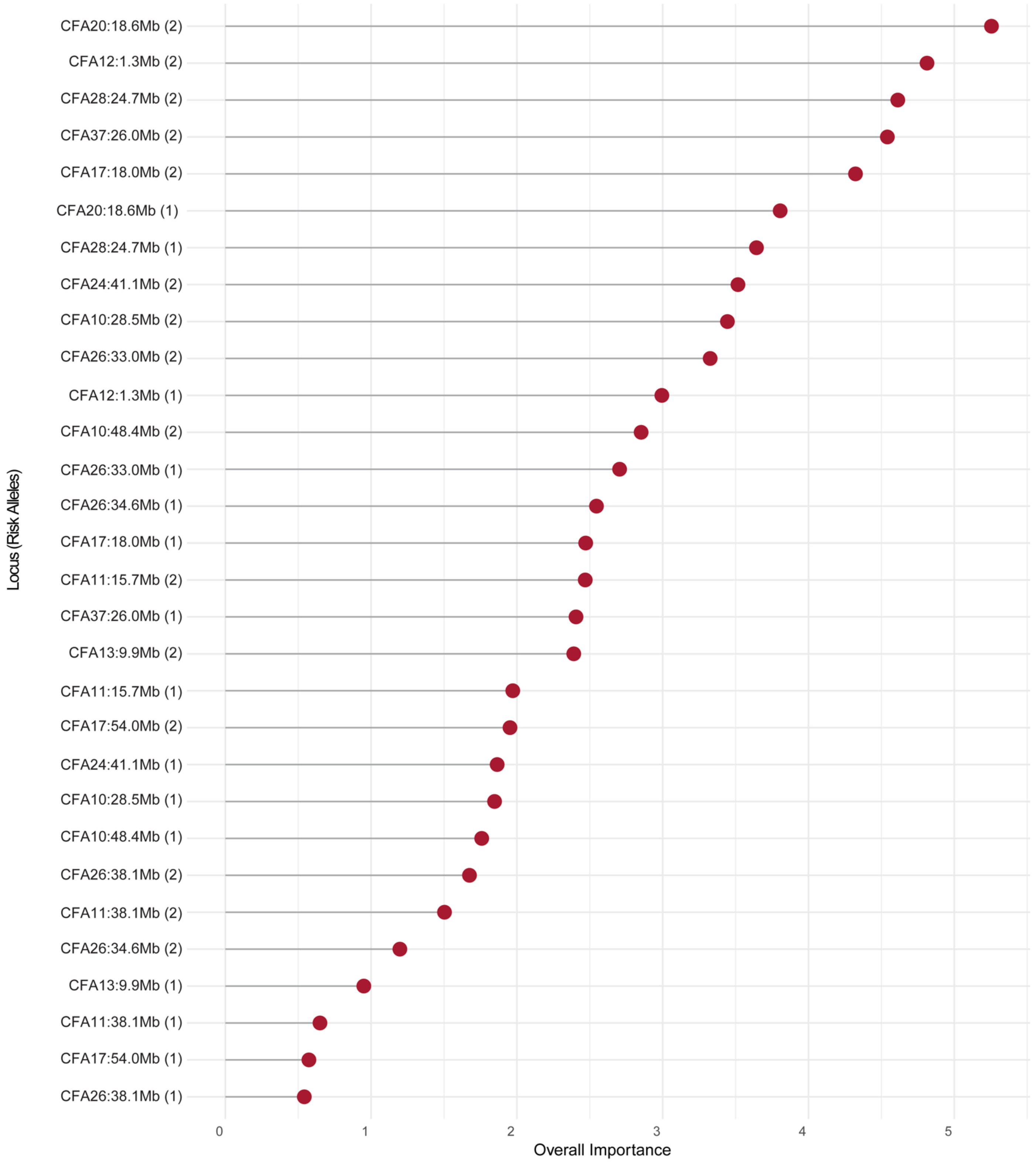


Figure S3. Importance of number of risk alleles at each locus in the 15-locus predictive model. The importance score (x-axis) ranks the relative contribution of one (1) or two (2) risk alleles at each locus (y-axis) to the model's predictive accuracy, with the highest score indicating the greatest effect within the model. The importance score is based on the absolute value of the t-statistic of each variable when applied to the model and was calculated using the varImp function in the caret R package. Importance is set to zero for zero copies of the risk allele.

A

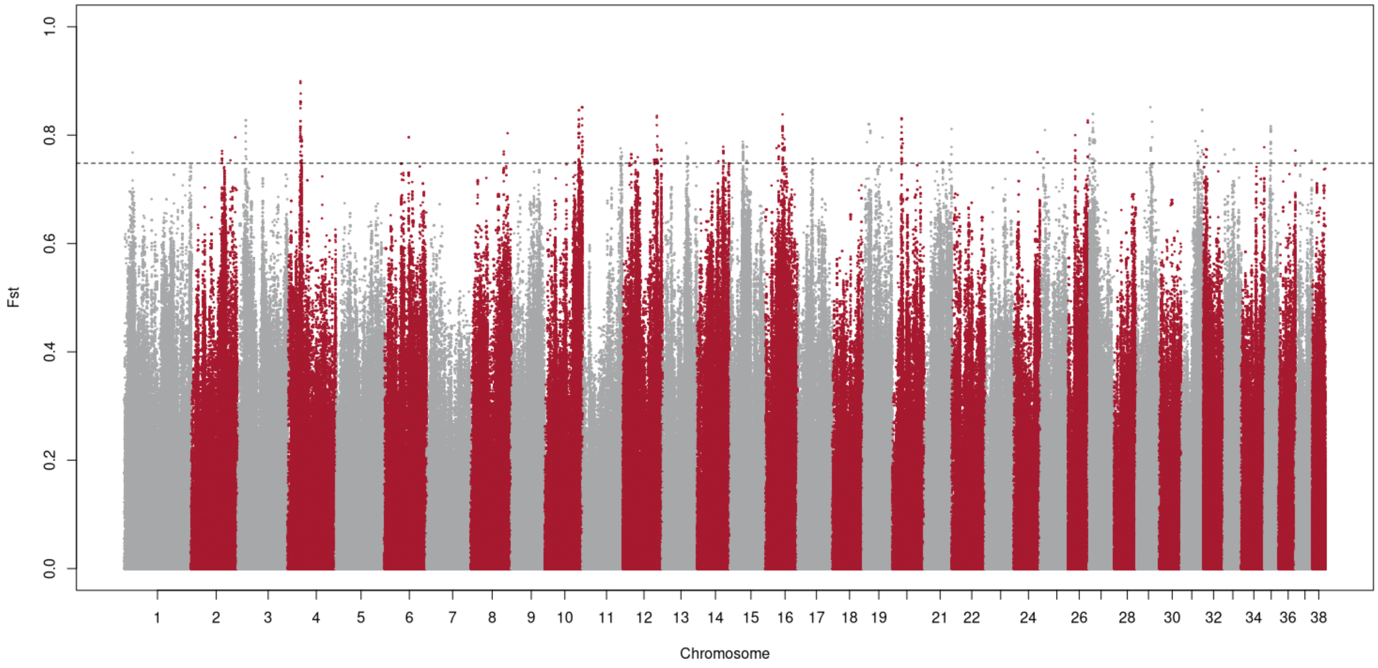


B

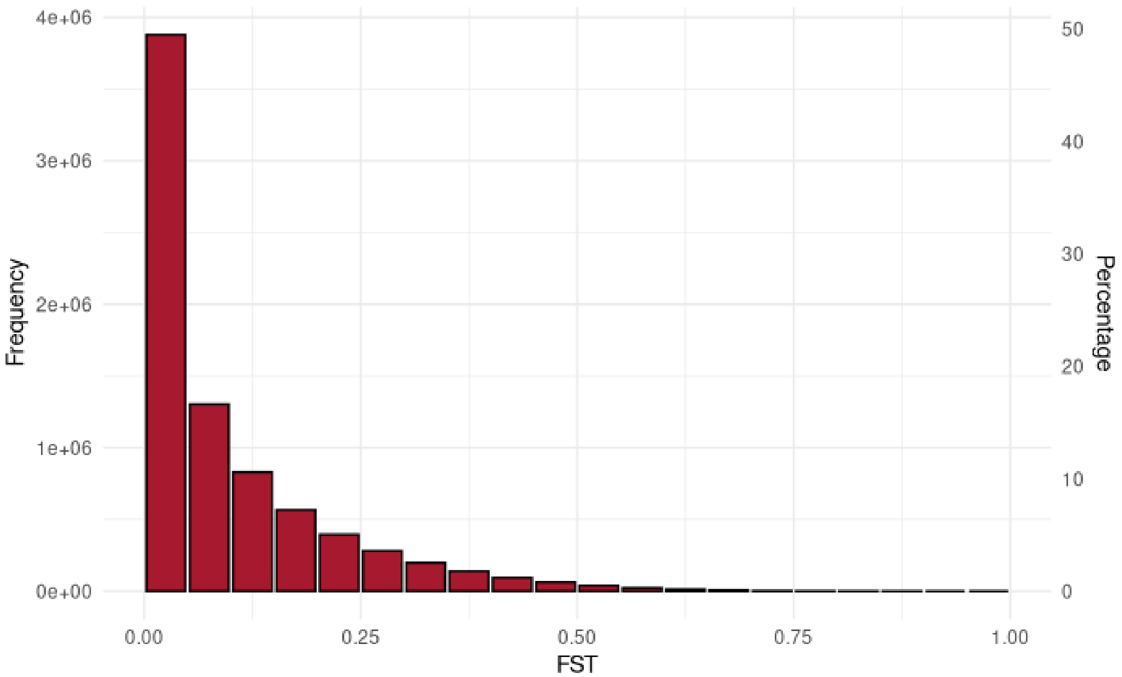


Figure S4. Across-breed fixation index (Fst) values. A) Fst values of 7,836,277 non-monomorphic variants comparing the Belgian Tervuren (n=69) and Belgian Sheepdogs (n=82) of unknown phenotype to Belgian Malinois (n=149) as reported by PLINK1.9. The dashed black line represents the 99.99% threshold. B) Distribution of Fst values; approximately half of all values lie between 0 and 0.05.

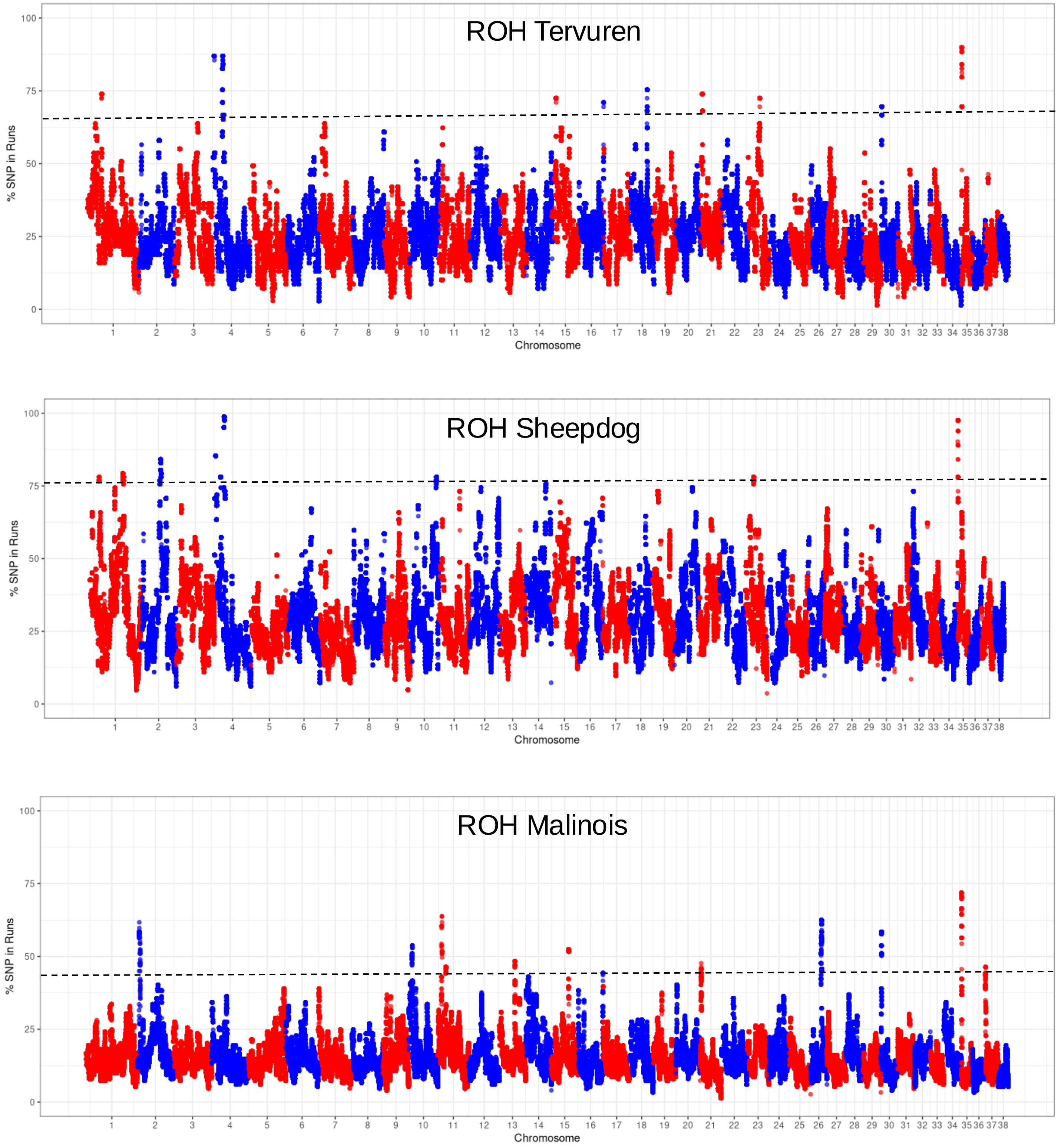


Figure S5. Incidence plots of runs of homozygosity (ROH) by breed. ROH were assessed in the Tervuren (n=69), Sheepdog (n=82), and Malinois (n=149) for 2,039,330 variants, with a minimum 500kb interval. The proportion of times each SNV is represented in an ROH within the population is shown. The dashed line represents the 99.9th percentile of runs calculated per breed.

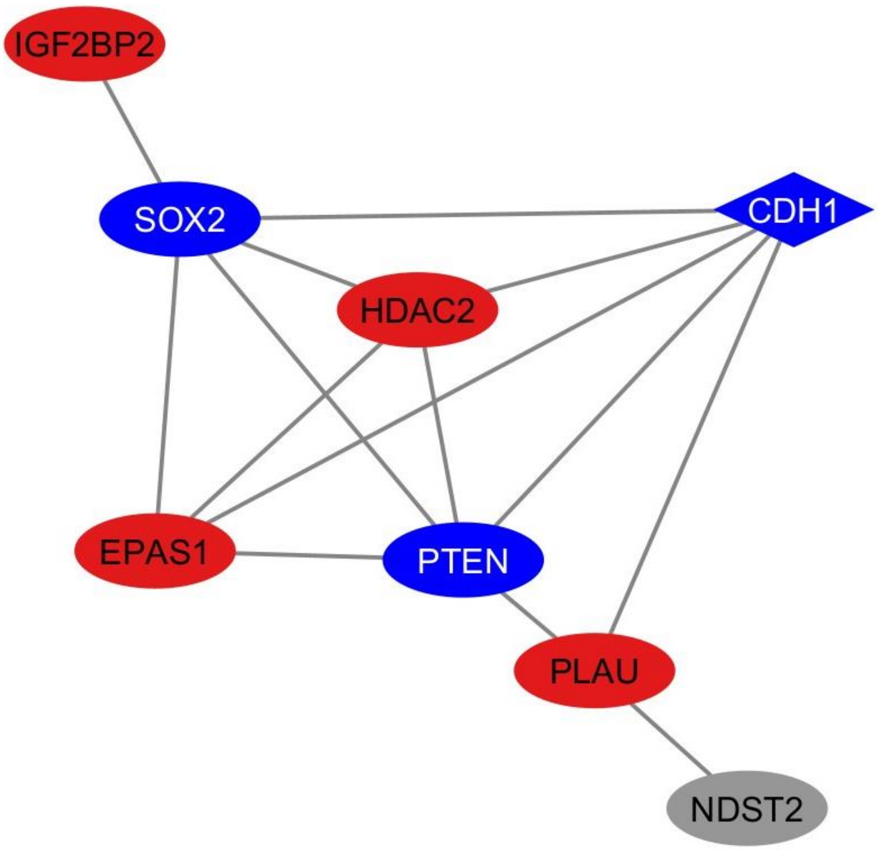


Figure S6. STRING protein network interactions. Proteins identified through GWAS and XP-nSL genomic analyses with reported interactions were included, as well as CDH1, whose expression is often lost in human diffuse GC. Protein interactions as defined by default STRING options are represented by gray lines. Blue shapes indicate downregulation and red shapes indicate upregulation of the protein or gene in human GC, while gray represents unknown. The figure was generated with Cytoscape.
